# assessPool: a flexible pipeline for population genomic analyses of pooled sequencing data

**DOI:** 10.1101/2024.10.09.617480

**Authors:** Evan B Freel, Emily E Conklin, Derek W Kraft, Jonathan L Whitney, Ingrid SS Knapp, Zac H Forsman, Robert J Toonen

## Abstract

Despite the dramatic decrease in high-throughput sequencing costs over time, sequencing the ideal number of individuals for population genetic inference remains prohibitively expensive. When research questions require only population-level resolution, pooling individual samples before sequencing (pool-seq) can substantially reduce costs while still providing allele frequencies of Single Nucleotide Polymorphisms (SNPs). However, analyzing pooled data is comparatively difficult and less standardized than individual-based analyses. Although several programs have been developed to handle pool-seq data, most require extensive formatting or programming skills to operate. Here we introduce assessPool, an open-source R and Bash pipeline for pool-seq analyses with a focus on population structure. AssessPool accepts a Variant-Call Format (VCF) file and a FASTA-formatted reference, providing a straightforward transition from commonly used pipelines such as Stacks or dDocent. AssessPool handles varying numbers of pools and utilizes PoPoolation2 to generate locus-by-locus pairwise *F_ST_* values and associated Fisher T-test values as measures of population structure. Starting with a VCF file containing all identified SNPs, assessPool facilitates several key functionalities for population genetic analyses: i) filtering SNPs based on adjustable criteria with parameter suggestions for pool-seq data, ii) organizing data structures for analysis based on input pools, iii) creating customizable run scripts for F*_ST_* calculations using PoPoolation2 and/or the {poolfstat} R package, for all pairwise comparisons, iv) calculating locus-specific *F_ST_* values using PoPoolation2 and/or {poolfstat}, v) importing *F_ST_* output into a format compatible with R, vi) producing population genomic summary statistics, and vii) generating interactive plots to visualize and explore data. A pooled dataset generated from wild populations is used here to showcase the features of the assessPool pipeline for population genomic analyses.

## Introduction

Concern about environmental issues and the global decline in biodiversity have continued to grow over recent decades (Lubchenco, 1995; Waldron et al., 2017; Cowie, Bouchet & Fontaine, 2022). Many conservation issues including extinction risk, how and why populations evolve, loss of genetic diversity, ability to evolve in response to environmental changes, resolution of management units, fragmentation of populations and reduced connectivity can be addressed using genetic methods (Lande, 1988; Allendorf, Luikart & Aitken, 2013; Taylor, Dussex & van Heezik, 2017). High Throughput Sequencing (HTS) methods provide unprecedented insights by accessing vast quantities of genetic information, addressing complex issues with increasing speed and decreasing cost (Hedrick, 2001; Luikart et al., 2003; Mardis, 2008). As it becomes cheaper and easier to apply population genetic approaches at the genome scale the precision and accuracy of model parameters (such as effective population size and migration rates) can be greatly increased, broadening inferences and limiting locus specific biases (Luikart et al., 2003; Allendorf, Hohenlohe & Luikart, 2010; Toews & Brelsford, 2012; DiBattista et al., 2015; Teske et al., 2018).

While it is now possible to sequence complete genomes of many individuals, it is often still cost-prohibitive to employ individual-level sequencing for large wild non-model populations. Furthermore the insights gained by whole genome sequencing are not necessarily superior to some cheaper alternatives and are unlikely to result in improved conservation outcomes in many cases (Allendorf, Hohenlohe & Luikart, 2010; Bowen et al., 2014; McKinney et al., 2017; Meek & Larson, 2019). Due to trade-offs between cost and power in experimental design, reduced representation genomic sequencing methods, such as sequence capture (Grover, Salmon & Wendel, 2012; Schwarze et al., 2018) or restriction site-associated DNA (RAD) sequencing (Baird et al., 2008; Hohenlohe et al., 2010; Etter et al., 2012) have quickly become the most popular genetic techniques used in the current literature (Davey & Blaxter, 2010; Andrews & Luikart, 2014; Andrews et al., 2016; Harvey et al., 2016; Russello et al., 2020). While powerful and comparatively inexpensive, the cost and technical requirements for sequence capture methods or individual-based RAD-seq techniques are still out of reach for many researchers, especially in developing countries (Barber et al., 2014; Wepfer et al., 2021). Pooled sequencing (pool-seq) offers a powerful and cost-effective solution for population genomic studies based on allele frequencies across broad geographic areas. By pooling samples, researchers can obtain genomic-level data at a fraction of the cost compared to individual genotype-based or single locus studies (Kurland et al., 2019). While the number of individuals in a pool must be sufficiently large and care must be taken to ensure equimolar pooling (Anderson, Skaug & Barshis, 2014; Schlotterer et al., 2014; Anand et al., 2016), the accuracy of pooled approaches for estimating allele frequencies and uncovering fine-scale population structure has been validated in diverse taxa, including plants (Rellstab et al., 2013), fruit flies (Kofler, Betancourt & Schlotterer, 2012), and sharks (Kraft et al., 2020). However, a primary obstacle to this work is the lack of user-friendly analytical tools designed for analyzing pooled samples.

Currently, analyzing pooled data remains computationally challenging and unstandardized. Several programs have been developed specifically to handle pool-seq data; unfortunately, most still require specific formatting and programming skills. Our pipeline, assessPool, uses Bash and R code to process a file of identified variants, primarily Single Nucleotide Polymorphisms (SNPs), provided in Variant Call Format (.VCF) files. It expands on a series of bioinformatic tools to identify regions and magnitude of allele frequency differentiation across pools. Specifically, PoPoolation2 (Kofler, Pandey & Schlotterer, 2011) requires a separate Perl command and .sync file for each pairwise pool comparison to calculate pairwise *F_ST_*. Next, assessPool creates these necessary sync files and generates a shell script to run all desired pairwise comparisons. It allows for parameter adjustments at each step and executes the command in parallel (multithreading support using GNU Parallel (Tange, 2011)), thereby efficiently utilizing available computational resources.

Throughout the pipeline, visualizations are generated to help determine the impacts of specific filter thresholds and analytical parameters. These visualizations are critical for users to explore individual datasets and avoid scoring biases (Etter et al., 2012; Davey et al., 2013; DaCosta & Sorenson, 2014; Shafer et al., 2017; O’Leary et al., 2018). This ability to explore parameters allows the user to make informed decisions regarding complex choices that can impact the outcomes and conclusions of their studies. AssessPool then generates publication-ready visualizations, with both static and interactive visualizations that can be used without requiring extensive experience in population genetics or bioinformatics. In both SNP filtration and *F_ST_* calculation, small changes to parameters can greatly change downstream results (Shafer et al. 2017; O’Leary et al. 2018). Therefore, it is critically important to understand how each step in the process impacts an analysis and assessPool provides the visualization tools required to achieve this goal.

The assessPool pipeline operates on the downstream (post variant-calling) portion of data analysis, an area that was lacking for pool-seq data with existing tools. While programs do exist for calculating genetic differentiation from allele frequencies, such as PoPoolation2 (Kofler, Pandey & Schlotterer, 2011) and {poolfstat} (Hivert et al., 2018), the use of these programs often requires significant data manipulation per analysis. AssessPool provides the first user-friendly interface to largely automate this process, from a VCF file through data visualization.

## Materials and Methods

The required input for assessPool is a Variant Call Format (.VCF) file created using any of the standard practices in the field (e.g. FreeBayes (Garrison & Marth, 2012), GATK (McKenna et al., 2010) or Geneious (Drummond et al., 2010)). Here, data used to generate the visualization examples come from samples of the scleractinian coral *Montipora capitata*, collected from 10 islands in the Hawaiian archipelago (Hawaii’s Division of Natural Resources Special Activity Permits SAP-2006-02, SAP- 2008-99, SAP-2011-1, and SAP-2012-63), with each pool consisting of 24-57 individual samples. Libraries were prepared using the ezRAD protocol, following the steps outlined for equimolar pooling of individual samples (Toonen et al., 2013; Knapp et al., 2017) We use an input file with a reduced subset of approximately two million loci (before filtering) was used to efficiently run the example dataset for benchmarking and troubleshooting purposes. To generate the example VCF file we: i) trimmed raw reads via Trim Galore! (Krueger, 2012), ii) performed a *de novo* reference genome assembly via SPAdes (Bankevich et al., 2012), iii) mapped reads via bwa (Li & Durbin, 2009), and iv) performed variant calling via FreeBayes (Garrison & Marth, 2012). Mapping and variant calling were visualized using Tablet (Milne et al., 2013). Using this workflow (**Figure 1**) a VCF output from FreeBayes was used as input into assessPool to begin variant filtering and analysis.

**Figure 1.**
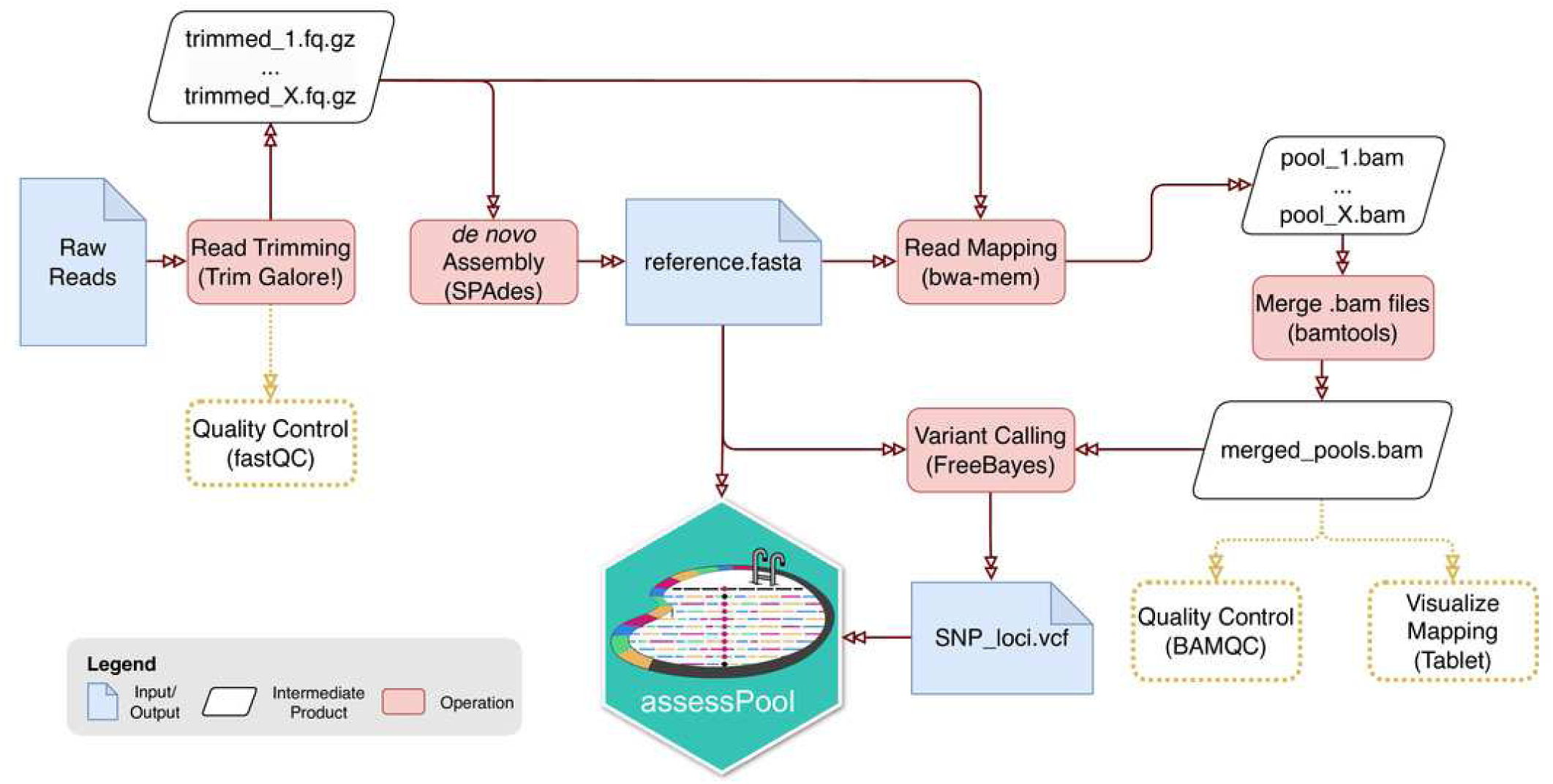
Schematic of the bioinformatic workflow from raw reads to the necessary assessPool input files. Blue polygons represent input and output files, white polygons are intermediate/temporary files, and red polygons notate general operational steps in the workflow. Dashed, yellow lines indicate optional quality control output files generated.

The assessPool pipeline is available on the ToBoDev GitHub repository, located at https://github.com/ToBoDev/assessPool and is run entirely from one R markdown document, using the interactive RStudio interface (RStudio Team, 2020). Both R and Bash shell code are implemented throughout and run on Unix-based operating systems.

Like many other bioinformatic pipelines, assessPool is dependent on a variety of specific tools to execute desired tasks at each step, with customization possible in a user-friendly format. This allows non-experts to efficiently alter analysis parameters from default pool-seq-specific recommendations without needing to know program-specific syntax.

An overview of the pipeline structure is provided in **Figure 2**. The required input files for assessPool are a VCF file generated by variant callers such as FreeBayes (Garrison & Marth, 2012), GATK (McKenna et al., 2010) or Geneious (Drummond et al., 2010), along with the corresponding FASTA format reference file used for mapping and variant calling. Example data are included in the GitHub repository (via Zenodo 10.5281/zenodo.13388658). It is important that the VCF file is appropriately separated by population (i.e., a genotype frequency column for each population at each locus in one file), because this structured approach lays the foundation for importing the VCF data into an R-compatible format for subsequent pipeline operations. In an R Markdown document, all dependencies are loaded via the R package {revn} and Bash installation scripts during initialization. Input files are then selected via an optional interactive prompt using {rstudio api}. Variants are then filtered through a series of recommended pool-seq specific filters using vcftools (Danecek et al., 2011) and vcflib (Garrison, 2012). Filters available are outlined in Table 1 and include minimum pool number, minimum quality score, minimum depth threshold across all pools, maximum missing genotypes (dropped due to low coverage), maximum allele length, mispaired reads, quality to depth ratio to remove highly undersplit loci, mean depth per site vs. quality score, and maximum mean depth to remove paralogs, and multicopy loci. The filtering process utilizes novel shell scripts that leverage GNU parallel (Tange, 2011) for parallel processing, greatly reducing filtering time with ample processing power and disc read/write speeds. The number of SNPs at each filter step is then output as an interactive HTML object via the R package {plotly} (Sievert, 2020), allowing the user to fine-tune filter parameters and locate potential issues before moving forward (**Figure 3 Error! Reference source not found.**).

**Figure 2.**
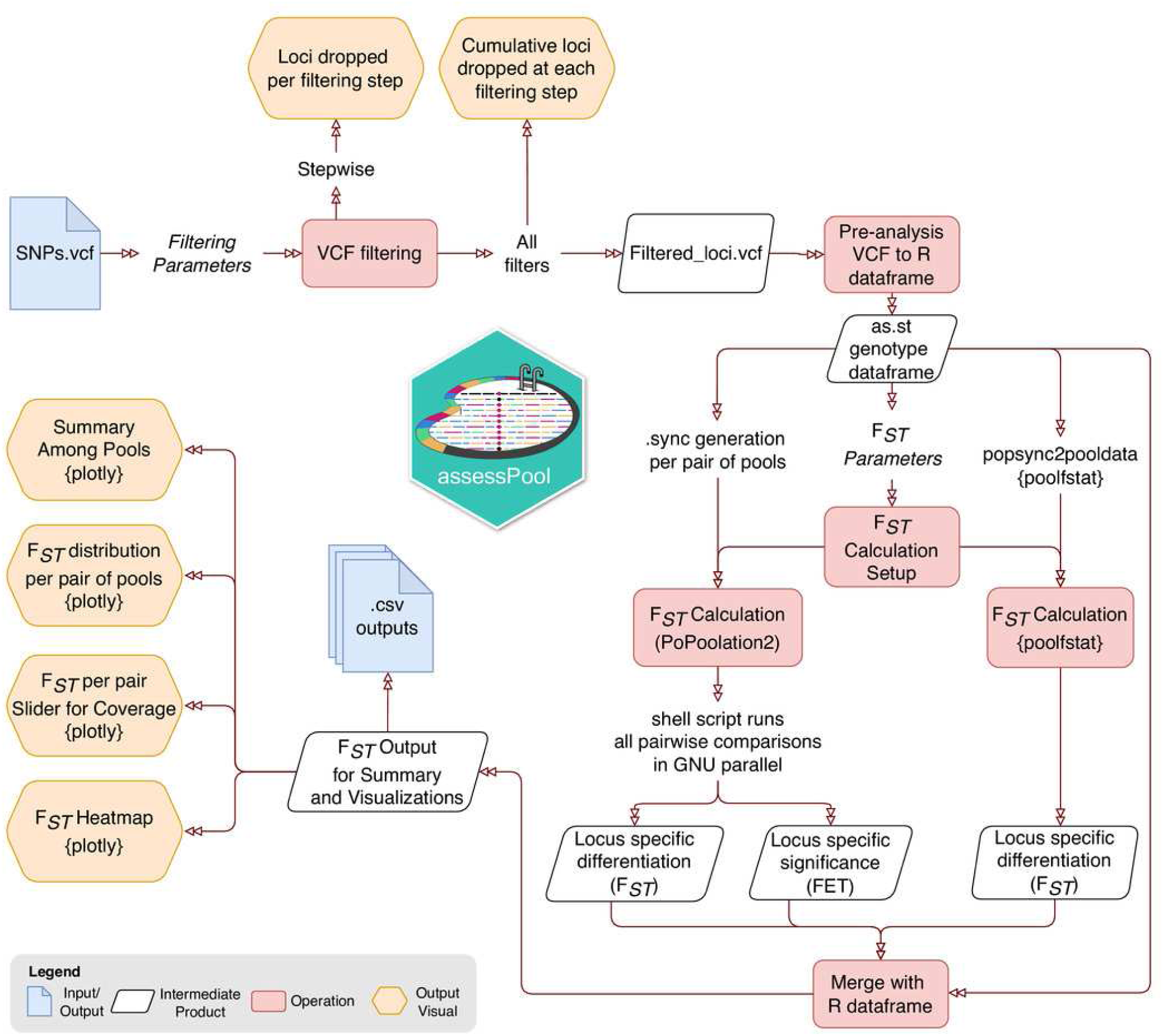
Schematic overview of assessPool modules and resulting visualizations. Blue polygons represent input and output files, white polygons are intermediary/temporary files, red polygons are operations steps and modules, and yellow polygons represent output visualizations.

**Figure 3.**
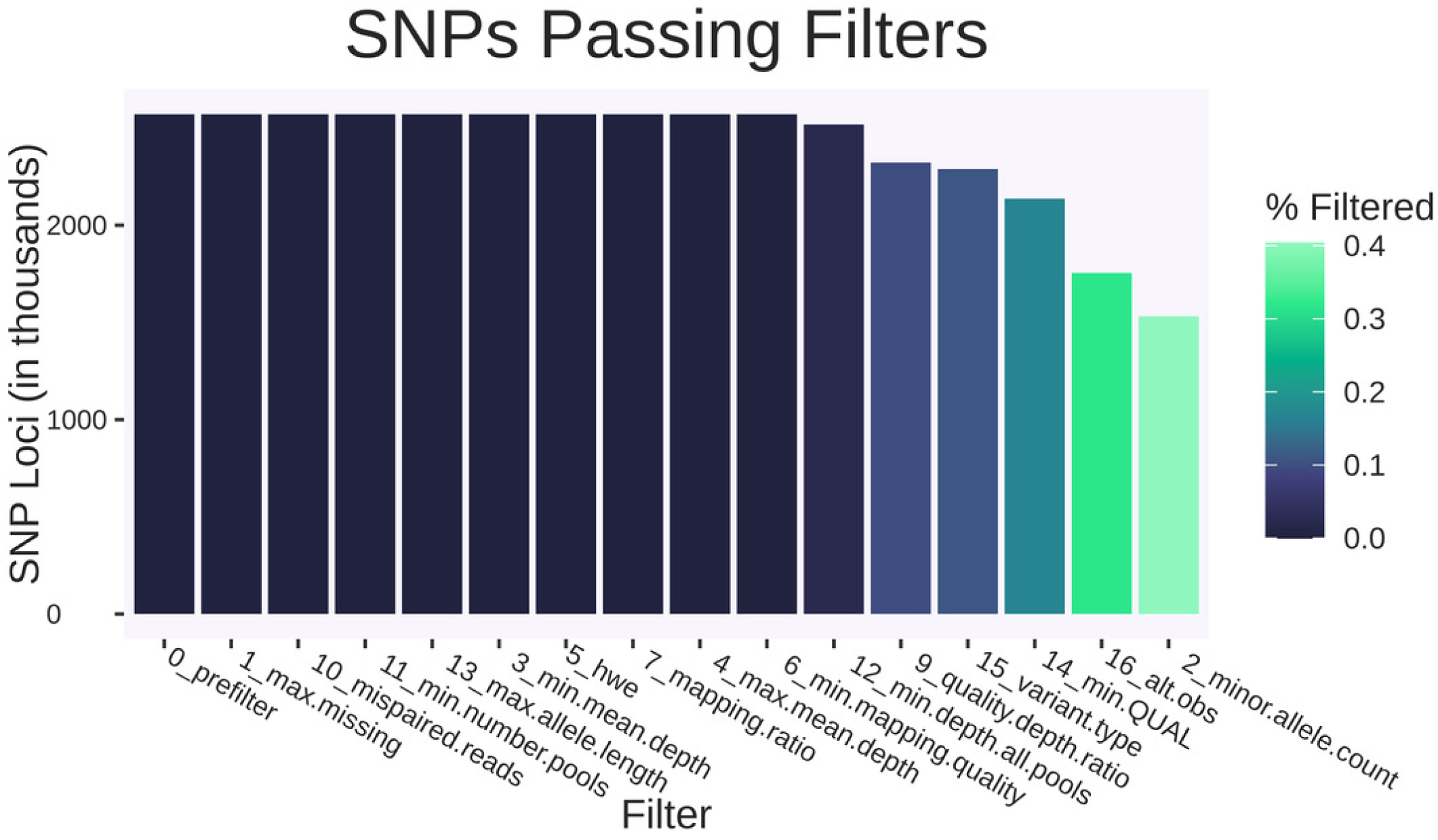
Static summary of SNP filtering at each individual filtering step. Shows the number of SNPs (in thousands) remaining from the input example example VCF file after applying each individual filtering step at default values. This can be used to determine where filtering steps drop data. In this example, many low-quality SNPs are conservatively filtered out. All filters are included after being run individually. Colored by proportion of loci filtered by each individual filtering step. Ordered by number of SNP loci filtered out.

**Table 1.**
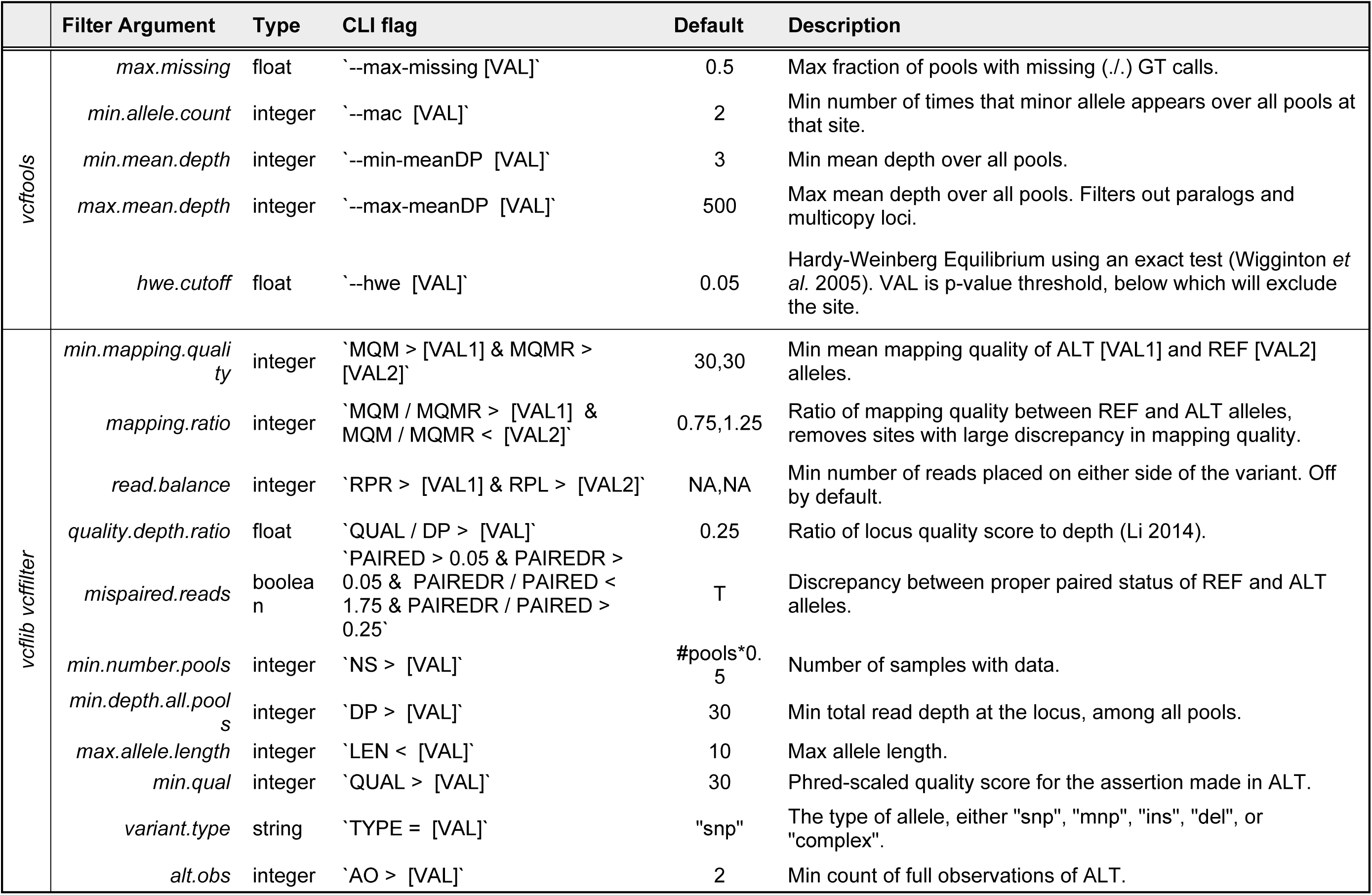
Filtering options available in assessPool for input Variant Call Format (VCF) file.

After VCF filtering, users select parameters for locus-specific pairwise *F_ST_* calculations using either PoPoolation2 (Kofler, Pandey & Schlotterer, 2011), the R package {poolfstat} (Hivert et al., 2018) or both. These *F_ST_* calculations are executed simultaneously and output to the console, producing essential summary information such as SNP counts and the location of exported variant loci. Following the *F_ST_* calculations, outputs from PoPoolation2 and/or {poolfstat} are converted into an R-compatible format to facilitate data visualization. During the development of assessPool, a primary objective was to create outputs that were accessible and informative, which led to the creation of interactive and static plots presenting summary statistics in several formats.

## Results

The assessPool filtering module can independently quantify the data filtered at each step utilizing vcftools and vcflib with multithreading support to generate a filtered VCF file. The first visualization output from the filtering module illustrates loci that pass filters when applied individually, aiding in the selection of filter parameters (**Figure 3**). After desired filtration parameters are selected, a final filtering summary output is generated to show number of cumulative sites lost during the VCF filtering module and (if selected) how many loci remain after thinning the VCF to avoid linked loci (**Figure 4**). After all loci-specific filters are completed, a final thinning step is available in order to ensure SNPs are less impacted by linkage and have are a minimum distance (default 300bp) apart. After completing F*_ST_* calculations and merging the results back into the R environment, summary statistic plots are output into a unified HMTL object. This object features a button-based interface allowing users to switch between plots showing SNP number, contig number, SNPs per contig, mean and standard deviation of *F_ST_* for all loci, and loci shared across all pools (**Figure 5** Error! Reference **source not found.**).

**Figure 4.**
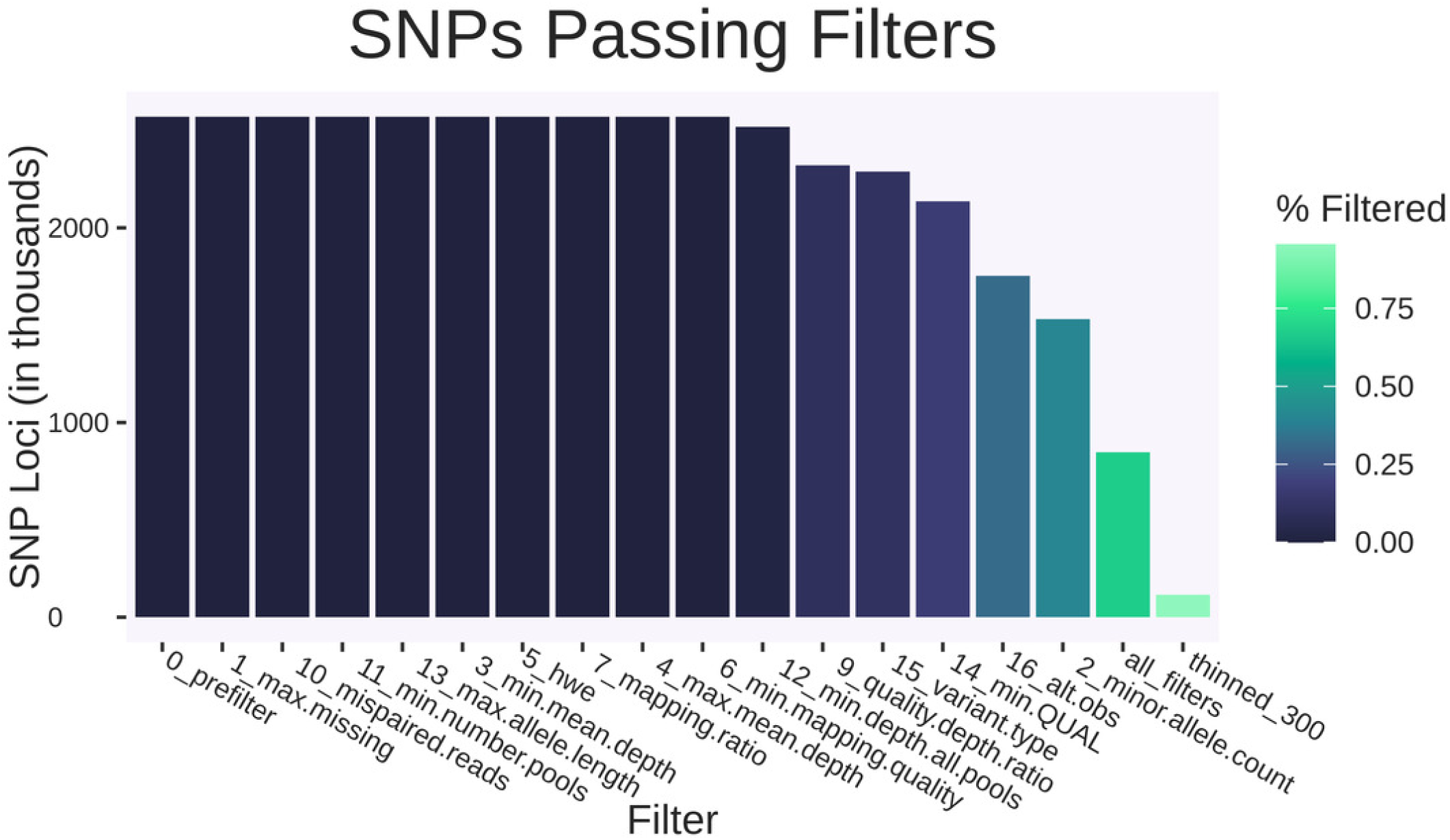
Static summary of cumulative SNP filtering at each successive step. Shows the cumulative number of SNPs (in thousands) remaining from the input example VCF file after applying each successive filtering step. In this example, many SNPs are conservatively filtered out with low quality. All filters are run consecutively, through an additional thinning step. Colored by proportion of loci filtered by consecutive filtering steps. Ordered by number of SNP loci filtered out.

**Figure 5.**
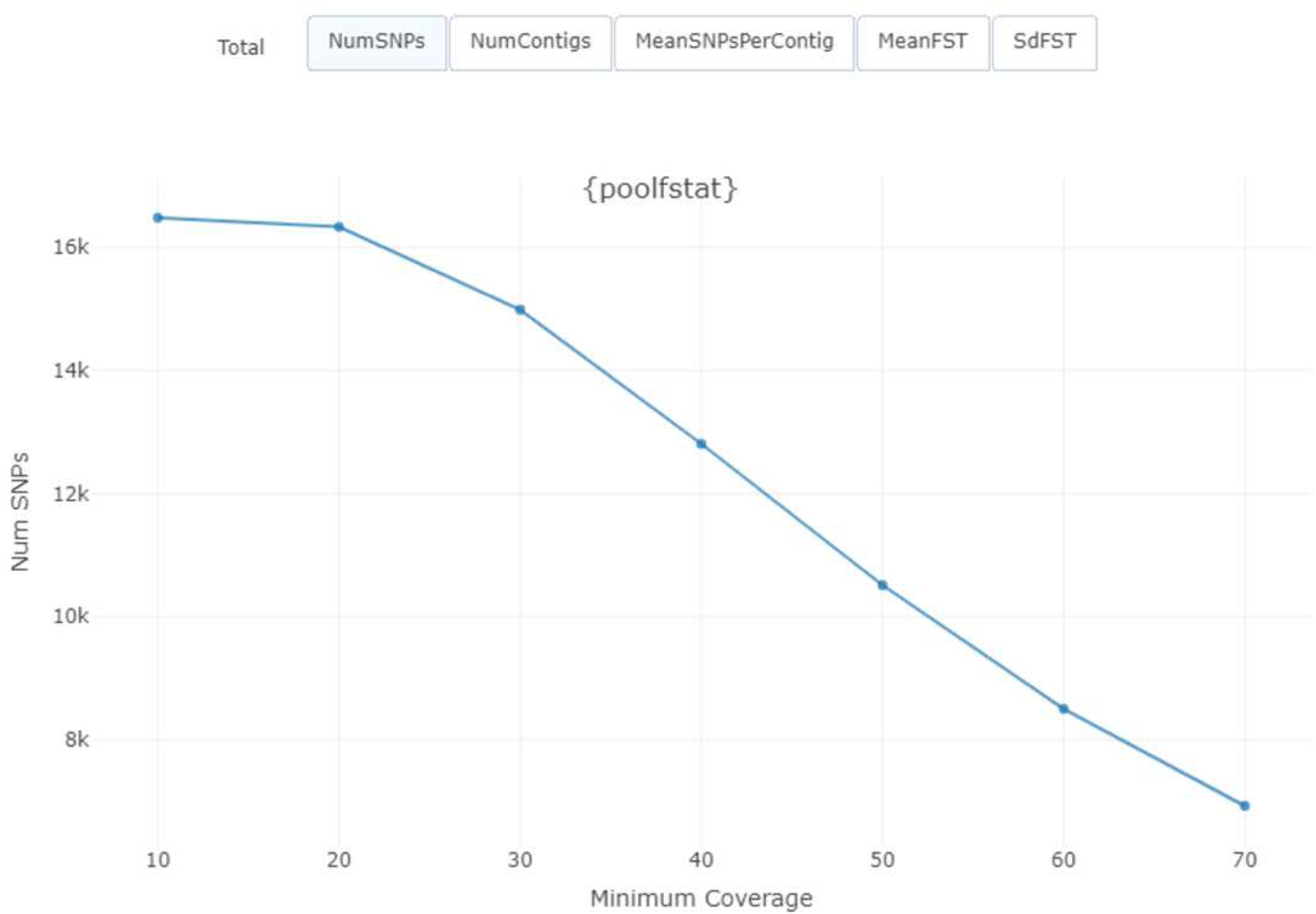
Interactive {plotly} widget of summary statistics among pools. Used to explore simple summary statistics in relation to minimum coverage cutofi. Tabs above the plot display minimum coverage vs the tab9s label and scale axes accordingly.

To simplify determination of coverage cutoffs, pairwise mean *F_ST_* is visualized in an interactive scatterplot. A slider allows exploration of how coverage influences both mean *F_ST_* and the number of SNPs included in each pairwise mean (**Figure 6**) **Error! Reference source not found.**. To visualize the distribution of each pairwise comparison, a boxplot object is produced which also has a slider for minimum coverage (**Figure 7**) **Error! Reference source not found.**. Hovertext highlights minimum, maximum, and quartiles for F_ST_. The user can then make an informed decision on the appropriate coverage cutoff to output as a static plot. Once an appropriate minimum coverage cutoff is determined, interactive heatmaps are produced (**Figures 8, 9**). Each pool is compared in pairwise fashion and the heatmap format highlights patterns of genomic differentiation. Heatmaps are generated for pairwise F*_ST_*, mean coverage, number of SNP loci, and significance. Significance is determined using a one-sided t-test to determine if all of the loci in a given pairwise comparison differs significantly from F*_ST_* = 0.

**Figure 6.**
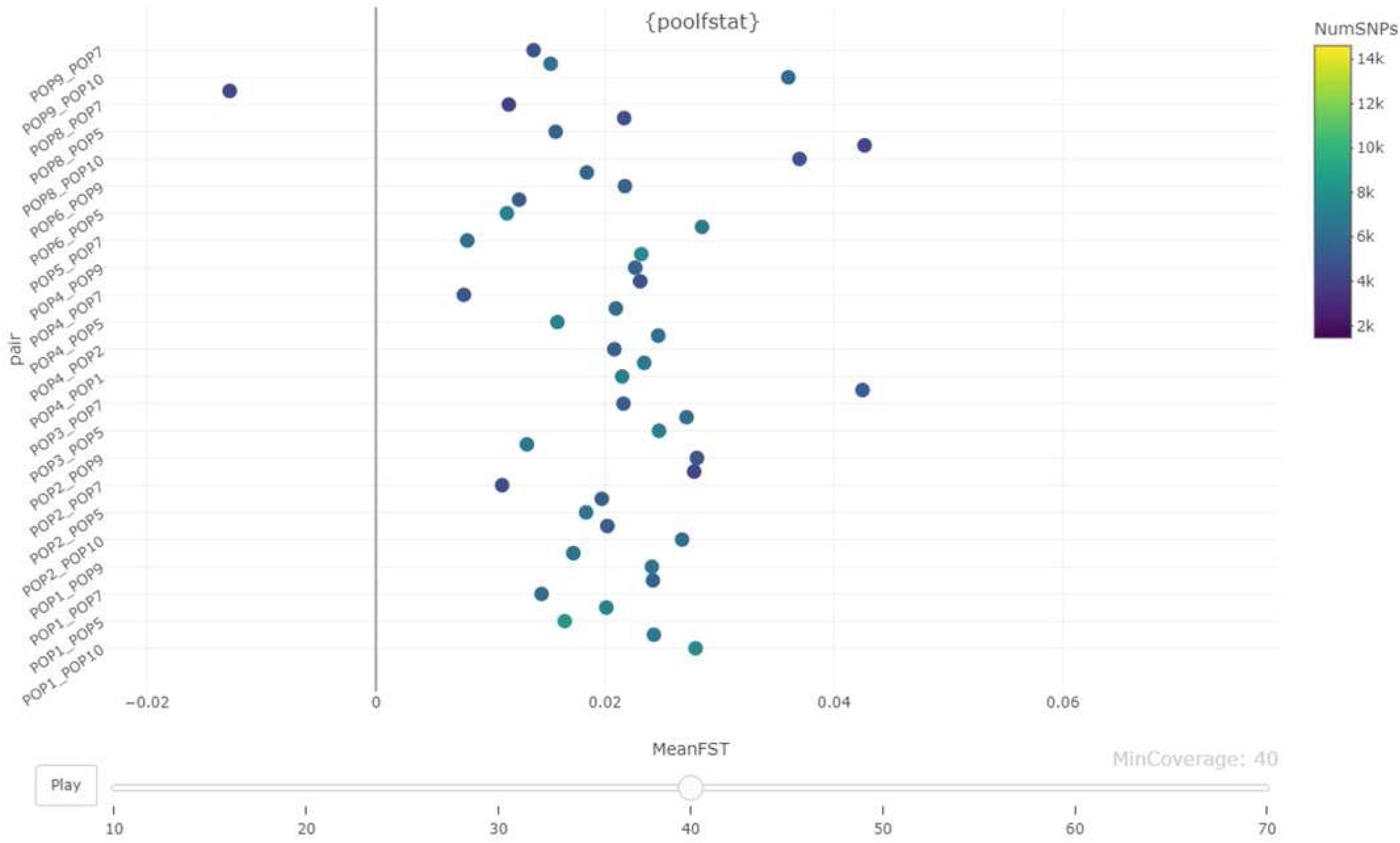
Interactive {plotly} scatterplot of mean F*_ST_* per pair of populations, adjustable by minimum coverage per locus. The slider on the bottom adjusts minimum coverage to visualize impact on broader pairwise F*_ST_* patterns and number of included SNPs. Colored by number of SNP loci.

**Figure 7.**
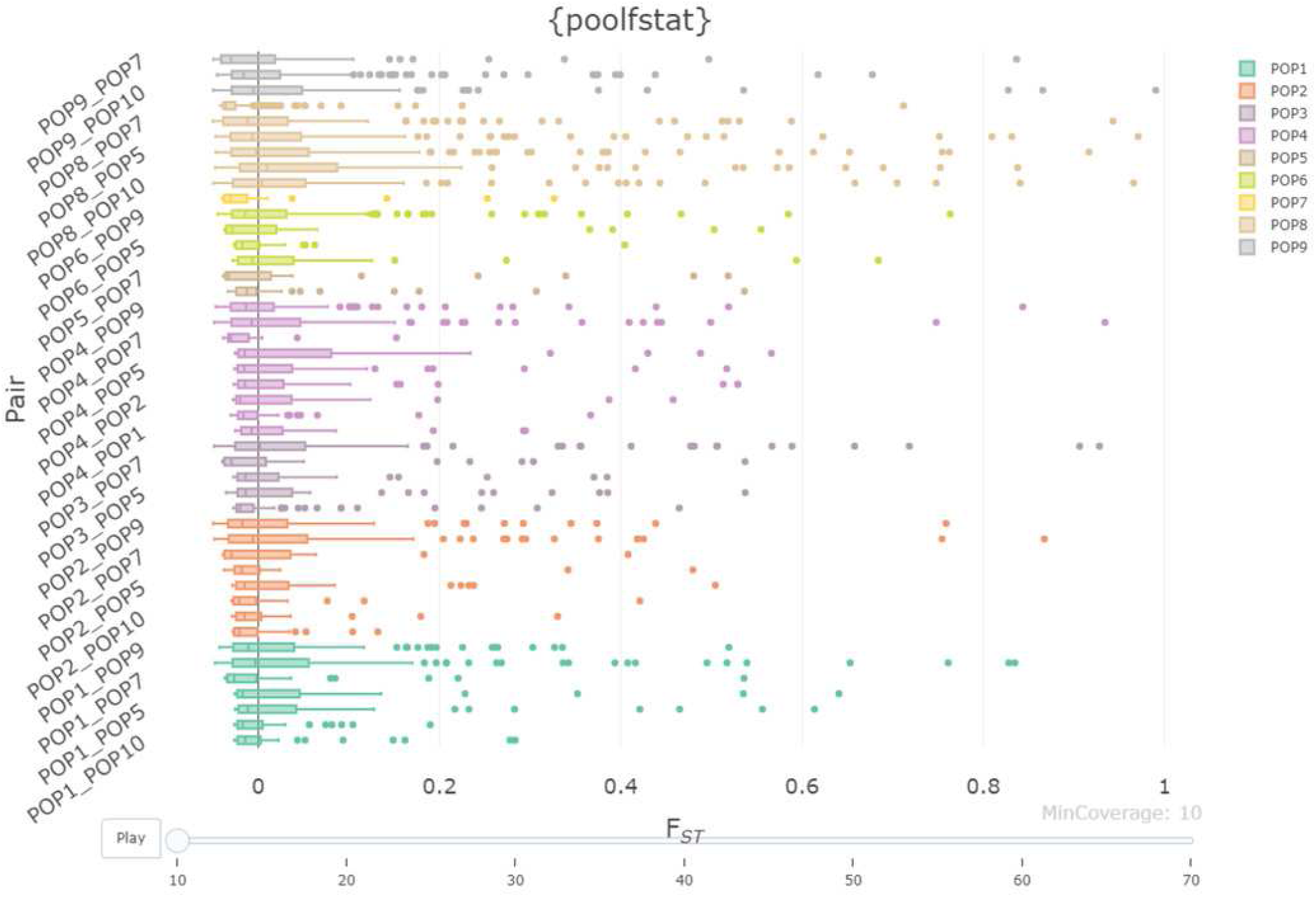
Interactive {plotly} boxplot of F*_ST_* distribution pair of populations, adjustable by minimum coverage per locus. Colored by pool (inherited from the left side of a given 8POPX_POPY9 pair of pools). The slider on the bottom can be used to adjust minimum coverage.

**Figure 8.**
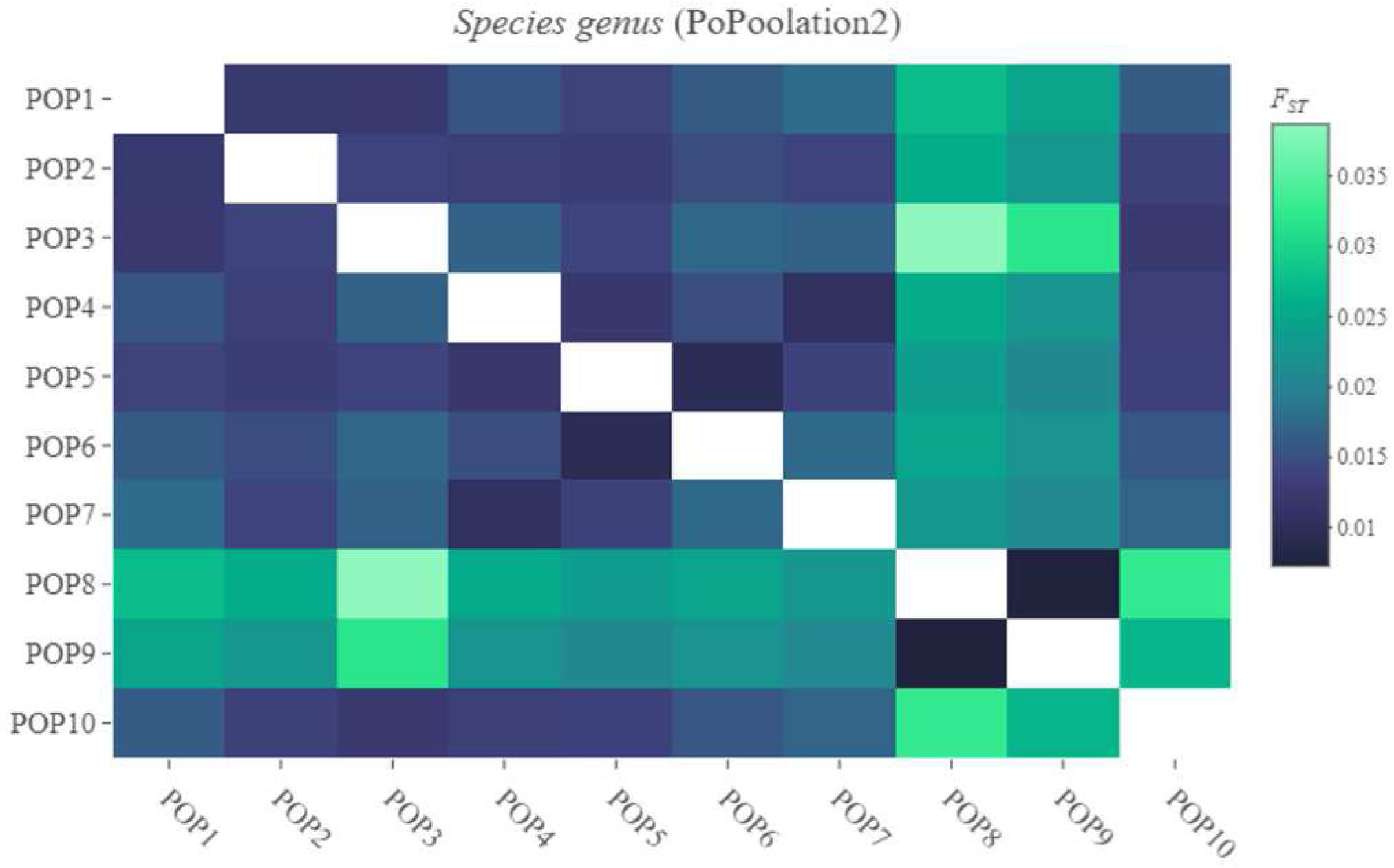
Interactive {plotly} heatmap of mean F*_ST_* for each pairwise comparison. Hovertext on each cell of the heatmap displays the exact F*_ST_* value for the pairwise comparison selected. Species name and package used for F*_ST_*calculation displayed as title.

**Figure 9.**
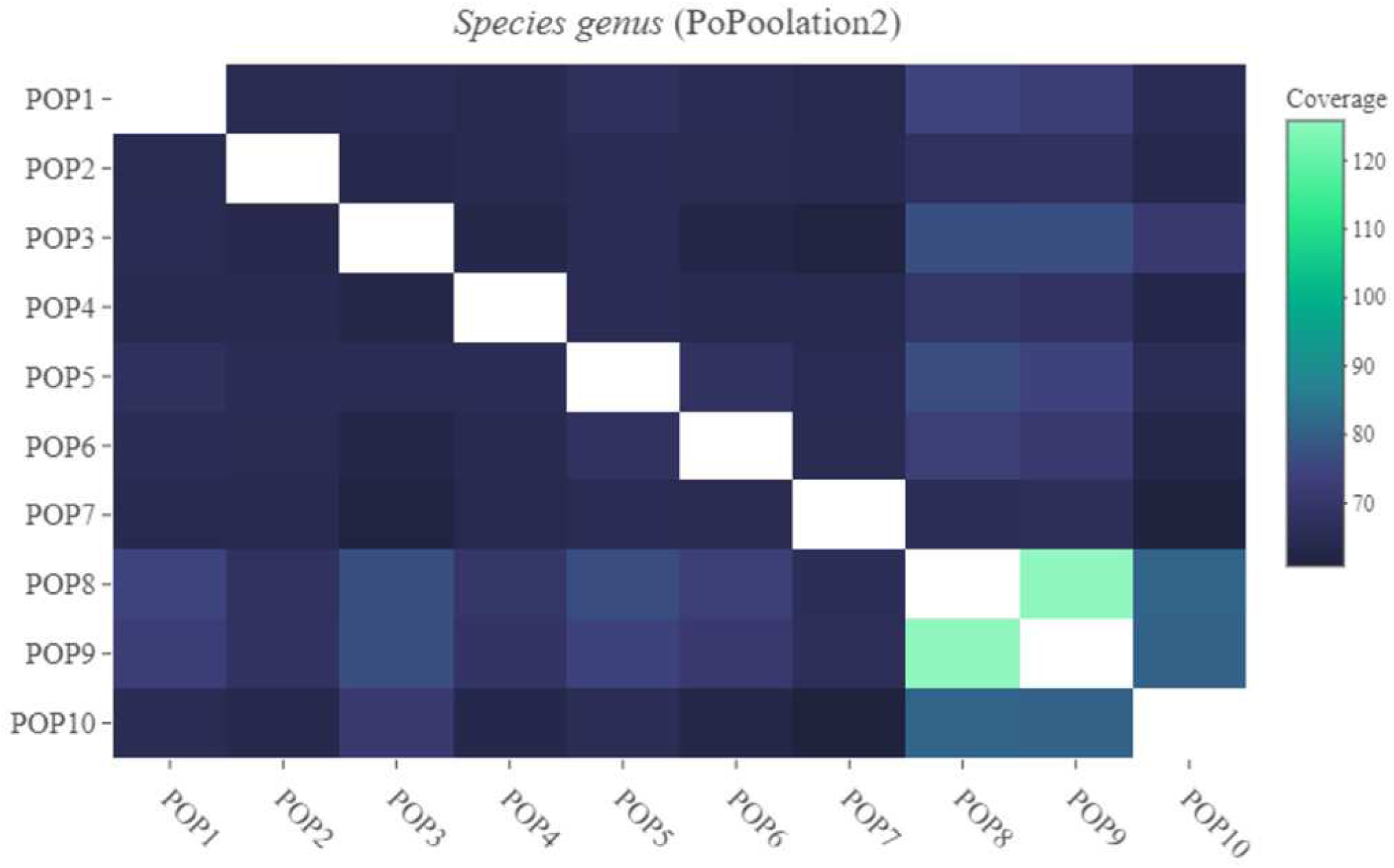
Interactive {plotly} heatmap of mean coverage for each pairwise comparison. Hovertext on each cell of the heatmap displays exact mean coverage for the pairwise comparison selected. Species name and package used for F*_ST_* calculation displayed as title.

## Discussion

AssessPool streamlines pool-seq population genomic analyses with broad applicability. This pipeline minimizes the need for developing custom analysis workflows to calculate locus-specific differentiation and visualize pool-seq data after variants are identified. Such standardized and reproducible workflows promote consistency and comparability across datasets and studies. Here we highlighted the tool’s utility and provided some applications for researchers in need of effective pool-seq analysis tools.

While several programs for handling pool-seq data already exist, assessPool expands on these tools by simplifying analyses and visualizing results. In SNP filtration and *F_ST_* calculation, small changes to parameters can change downstream results (Shafer et al., 2017; O’Leary et al., 2018). AssessPool’s interactive visualizations help users evaluate how their chosen filter thresholds and analysis parameters might influence their results and conclusions. This tool allows users to explore parameter space to better understand and optimize their analyses facilitating informed decisions regarding complex and interacting parameter choices. By producing publication-ready visualizations without requiring extensive expertise in population genetics or bioinformatics, assessPool streamlines the process of data exploration to publication.

Population genetic analyses are often based on allele frequency estimates within and among populations. While individuals are typically sequenced or genotyped separately, these analyses, such as *F_ST_* calculations, often combine individual data to calculate allele frequencies for a given location (Weir & Hill, 2002; Meirmans, 2006; Bird et al., 2011). Pool-seq, despite its drawbacks in precluding individual genotype-based analyses (Porras-Hurtado et al., 2013), such as *STRUCTURE* (Pritchard, Stephens & Donnelly, 2000; Falush, Stephens & Pritchard, 2007; Hubisz et al., 2009) proves advantageous for broad population-level comparisons based on allele frequencies. This approach avoids the unnecessary cost and potential waste associated with individual sequencing, making population genomic analyses accessible to more research groups. For many research questions, pooling samples provides cost-effective, accurate allele frequency estimates across many thousands of loci within population for less cost than running a single mitochondrial locus in some cases (Kraft et al., 2020). This cost reduction comes at a time when emerging, and expensive technologies may deter resource-limited research groups from advancing their research goals through pool-seq.

With assessPool, we provide a promising tool for conservation genetic applications by streamlining analysis of existing pool-seq population genetic approaches. Pool-seq applications can range from resolving fine scale genetic structure at small spatial scales, to delineating global stock structure of fisheries species. In each case, assessPool standardizes and optimizes the downstream analysis of pool-seq data. For example, aided by an assessPool-based workflow, Kraft et al. (2020) discovered distinct stock structure of silky sharks (*Cacharhinus falciformis*), previously undetected using mitochondrial markers alone. The pipeline facilitated automated handling and data formatting for pairwise comparisons of silky sharks from four spatially distinct populations (Gulf of Mexico, Northern Atlantic, Brazil, and Red Sea).

Likewise, pool-seq provides a relatively quick and cost-effective method for identifying outlier loci that are highly differentiated among groups. The move towards genomic approaches has advanced the field by enabling the scanning of larger portions of the genome to uncover patterns of genetic differentiation. For researchers utilizing pool-seq approaches, the pipeline presented here provides a standardized, and informed workflow for downstream analysis of pool-seq data. Besides aiding in management decisions, assessPool also holds potential for industry applications that often look to genome-wide association studies (Sharma et al., 2015; Visscher et al., 2017). Decreasing sequencing costs has made pool-seq an attractive option for identifying outliers as candidate loci for further research (Kofler, Betancourt & Schlotterer, 2012; Neethiraj et al., 2017; Taus, Futschik & Schlotterer, 2017).

Preliminary pool-seq analysis on groups of individuals expected to express genetic differentiation can be achieved far more quickly and affordably than with genome-wide association studies (Dennenmoser et al., 2017; Guirao-Rico & Gonzalez, 2021). The assessPool pipeline supports such research by streamlining data input from a raw VCF, through filtering and data preparation, for two frequently used pool-seq analysis tools, PoPoolation2 and {poolfstat}. The output from each is then conveniently organized for rapid data visualization in R. For those interested in exploring regions of high genetic differentiation, assessPool will also output all loci above a desired *F_ST_* value in FASTA format for straightforward downstream analysis needs (e.g. BLAST). For example, assessPool was used to streamline the identification of regions with significant allele frequency differentiation between freshwater and seawater reared lines of Tilapia (Freel, 2024). In essence, pool-seq allows the rapid and cost-effective genome scanning of individuals from differing environments (such as experimental treatments, rearing techniques, color morphs, or environmental tolerance groups) to identify regions of genetic differentiation for more targeted research. Our assessPool pipeline addresses the need for making pool-seq data more accessible and usable. Combined with the cost savings associated with population genetic approaches based on population allele frequencies, this should facilitate broader adoption of pool-seq in the field.

## Acknowledgements

We would like to express our gratitude to the many ToBo Lab members, particularly Richard Coleman, ‘Ale’alani Dudoit, and Cataixa Lopez, who used assessPool during development and helped identify issues and suggest key feature improvements. We would also like to thank Iliana Baums, Tanya Beirne, Dave Carlon, Greg Conception, Matt Craig, Jeff Eble, Scott Godwin, Matt Iacchei, Frederique Kandel, Steve Karl, Jim Maragos, Bob Moffitt, Joe O’Malley, Lawrie Provost, Jennifer Salerno, Derek Skillings, Michael Stat, Ben Wainwright, and Kim Weersing, for their efforts in sample collection.

---

Table 1

Filtering options available in assessPool for input Variant Call Format (VCF) file.

The table is split by which filtering program is used (vcftools or vcfiib). The equivalent Command Line Interface (CLI) fiag is shown with [VAL] used as a placeholder for assessPool’s default value of a given filter.

Manuscript to be reviewed

## References

1. Allendorf FW, Hohenlohe PA, Luikart G. 2010. Genomics and the future of conservation genetics. Nature Reviews Genetics 11:697–709. DOI: 10.1038/nrg2844.

2. Allendorf F, Luikart G, Aitken S. 2013. Conservation and the genetics of populations. Chichester, West Sussex: Wiley-Blackwell.

3. Anand S, Mangano E, Barizzone N, Bordoni R, Sorosina M, Clarelli F, Corrado L, Boneschi FM, D’Alfonso S, De Bellis G. 2016. Next generation sequencing of pooled samples: Guideline for variants’ filtering. Scientific Reports 6:1–9. DOI: 10.1038/srep33735.

4. Anderson EC, Skaug HJ, Barshis DJ. 2014. *Next-generation sequencing for molecular ecology: A caveat regarding pooled samples*. John Wiley & Sons, Ltd. DOI: 10.1111/mec.12609.

5. Andrews KR, Good JM, Miller MR, Luikart G, Hohenlohe PA. 2016. Harnessing the power of RADseq for ecological and evolutionary genomics. Nat Rev Genet advance on:81–92. DOI: 10.1038/nrg.2015.28.

6. Andrews KR, Luikart G. 2014. Recent novel approaches for population genomics data analysis. Molecular Ecology 23:1661–1667. DOI: 10.1111/mec.12686.

7. Baird NA, Etter PD, Atwood TS, Currey MC, Shiver AL, Lewis ZA, Selker EU, Cresko WA, Johnson EA. 2008. Rapid SNP Discovery and Genetic Mapping Using Sequenced RAD Markers. PLoS ONE 3:e3376. DOI: 10.1371/journal.pone.0003376.

8. Bankevich A, Nurk S, Antipov D, Gurevich AA, Dvorkin M, Kulikov AS, Lesin VM, Nikolenko SI, Pham S, Prjibelski AD, Pyshkin A V., Sirotkin A V., Vyahhi N, Tesler G, Alekseyev MA, Pevzner PA. 2012. SPAdes: A New Genome Assembly Algorithm and Its Applications to Single-Cell Sequencing. Journal of Computational Biology 19:455–477. DOI: 10.1089/cmb.2012.0021.

9. Barber PH, Carmen Ablan-Lagman MA, Berlinck RG, Cahyani D, Crandall ED, Ravago- Gotanco R, Antonette Juinio-Mefiez M, Ngurah Mahardika I, Shanker K, Starger CJ, Hamid Toha AA, Anggoro AW, Willette DA. 2014. Advancing biodiversity research in developing countries: the need for changing paradigms OA Open access content. Bull Mar Sci 90:187–210. DOI: 10.5343/bms.2012.1108.

10. Bird CE, Karl SA, Smouse PE, Toonen RJ. 2011. Detecting and measuring genetic differentiation. In: *Phylogeography and Population Genetics in Crustacea*. 1–55.

11. Bowen BW, Shanker K, Yasuda N, Celia M, Malay MC (Machel) D, von der Heyden S, Paulay G, Rocha LA, Selkoe KA, Barber PH, Williams ST, Lessios HA, Crandall ED, Bernardi G, Meyer CP, Carpenter KE, Toonen RJ. 2014. Phylogeography unplugged: comparative surveys in the genomic era. Bulletin of Marine Science 90:13-46. DOI: 10.5343/bms.2013.1007.

12. Cowie RH, Bouchet P, Fontaine B. 2022. The Sixth Mass Extinction: fact, fiction or speculation? Biological Reviews 97:640–663. DOI: 10.1111/BRV.12816.

13. DaCosta JM, Sorenson MD. 2014. Amplification Biases and Consistent Recovery of Loci in a Double-Digest RAD-seq Protocol. PLoS ONE 9:e106713. DOI: 10.1371/journal.pone.0106713.

14. Danecek P, Auton A, Abecasis G, Albers CA, Banks E, DePristo MA, Handsaker RE, Lunter G, Marth GT, Sherry ST, McVean G, Durbin R. 2011. The variant call format and VCFtools. Bioinformatics 27:2156–2158. DOI: 10.1093/bioinformatics/btr330.

15. Davey JW, Blaxter ML. 2010. RADSeq: next-generation population genetics. Briefings in Functional Genomics 9:416–423. DOI: 10.1093/bfgp/elq031.

16. Davey JW, Cezard T, Fuentes0Utrilla P, Eland C, Gharbi K, Blaxter ML. 2013. Special features of RAD Sequencing data: implications for genotyping. Molecular Ecology 22:3151-3164. DOI: 10.1111/mec.12084.

17. Dennenmoser S, Vamosi SM, Nolte AW, Rogers SM. 2017. Adaptive genomic divergence under high gene flow between freshwater and brackish-water ecotypes of prickly sculpin (Cottus asper) revealed by Pool-Seq. Molecular Ecology 26:25–42. DOI: 10.1111/mec.13805.

18. DiBattista JD, Waldrop E, Rocha LA, Craig MT, Berumen ML, Bowen BW. 2015. Blinded by the bright: a lack of congruence between colour morphs, phylogeography and taxonomy for a cosmopolitan Indo-Pacific butterflyfish, Chaetodon auriga. Journal of Biogeography 42:1919-1929. DOI: 10.1111/jbi.12572.

19. Drummond A, Ashton B, Buxton S, Cheung M, Cooper A, Heled J, Kearse M, Moir R, Stones-Havas S, Sturrock S, Thierer T, Wilson A. 2010. Geneious v6.1.8.

20. Etter PD, Bassham S, Hohenlohe PA, Johnson EA, Cresko WA. 2012. SNP Discovery and Genotyping for Evolutionary Genetics Using RAD Sequencing. In: Humana Press, 157-178. DOI: 10.1007/978-1-61779-228-1_9.

21. Falush D, Stephens M, Pritchard JK. 2007. Inference of population structure using multilocus genotype data: dominant markers and null alleles. Molecular Ecology Notes 7:574–578. DOI: 10.1111/j.1471-8286.2007.01758.x.

22. Freel EB. 2024. Population Genomic Tools and Applications of Pooled Sequencing Data. University of Hawai’i at Mānoa.

23. Garrison E. 2012. Vcflib: A C++ library for parsing and manipulating VCF files.

24. Garrison E, Marth G. 2012. Haplotype-based variant detection from short-read sequencing. *arXiv preprint* arXiv:1207.

25. Grover CE, Salmon A, Wendel JF. 2012. Targeted sequence capture as a powerful tool for evolutionary analysis. American Journal of Botany 99:312–319. DOI: 10.3732/ajb.1100323.

26. Guirao-Rico S, Gonzalez J. 2021. Benchmarking the performance of Pool-seq SNP callers using simulated and real sequencing data. Molecular Ecology Resources 21:1216–1229. DOI: 10.1111/1755-0998.13343.

27. Harvey MG, Smith BT, Glenn TC, Faircloth BC, Brumfield RT. 2016. Sequence Capture versus Restriction Site Associated DNA Sequencing for Shallow Systematics. Systematic Biology 65:910–924. DOI: 10.1093/sysbio/syw036.

28. Hedrick PW. 2001. Conservation genetics: where are we now? Trends in Ecology & Evolution 16:629–636. DOI: 10.1016/S0169-5347(01)02282-0.

29. Hivert V, Leblois R, Petit EJ, Gautier M, Vitalis R. 2018. Measuring Genetic Differentiation from Pool-seq Data. Genetics 210:315–330. DOI: 10.1534/genetics.118.300900.

30. Hohenlohe PA, Bassham S, Etter PD, Stiffler N, Johnson EA, Cresko WA. 2010. Population Genomics of Parallel Adaptation in Threespine Stickleback using Sequenced RAD Tags. PLoS Genetics 6:e1000862. DOI: 10.1371/journal.pgen.1000862.

31. Hubisz MJ, Falush D, Stephens M, Pritchard JK. 2009. Inferring weak population structure with the assistance of sample group information. Molecular Ecology Resources 9:1322–1332. DOI: 10.1111/j.1755-0998.2009.02591.x.

32. Knapp ISS, Puritz JB, Bird CE, Whitney JL, Sudek M, Forsman ZH, Toonen RJ. 2017. ezRAD- an accessible next-generation RAD sequencing protocol suitable for non- model organisms_v3.2. protocols.io. DOI: dx.doi.org/10.17504/protocols.io.e9pbh5n.

33. Kofler R, Betancourt AJ, Schlotterer C. 2012. Sequencing of Pooled DNA Samples (Pool-Seq) Uncovers Complex Dynamics of Transposable Element Insertions in Drosophila melanogaster. PLoS Genetics 8:e1002487. DOI: 10.1371/journal.pgen.1002487.

34. Kofler R, Pandey RV, Schlotterer C. 2011. PoPoolation2: identifying differentiation between populations using sequencing of pooled DNA samples (Pool-Seq). Bioinformatics 27:3435–3436. DOI: 10.1093/bioinformatics/btr589.

35. Kraft DW, Conklin EE, Barba EW, Hutchinson M, Toonen RJ, Forsman ZH, Bowen BW. 2020. Genomics versus mtDNA for resolving stock structure in the silky shark ( Carcharhinus falciformis ). PeerJ 8:e10186. DOI: 10.7717/peerj.10186.

36. Krueger F. 2012. Trim Galore: a wrapper tool around Cutadapt and FastQC to consistently apply quality and adapter trimming to FastQ files, with some extra functionality for MspI-digested RRBS-type (Reduced Representation Bisufite-Seq) libraries.

37. Kurland S, Wheat CW, Paz Celorio Mancera M, Kutschera VE, Hill J, Andersson A, Rubin C, Andersson L, Ryman N, Laikre L. 2019. Exploring a Pool0seq0only approach for gaining population genomic insights in nonmodel species. Ecology and Evolution 9:11448-11463. DOI: 10.1002/ece3.5646.

38. Lande R. 1988. Genetics and demography in biological conservation. Science 241:1455–1460. DOI: 10.1126/science.3420403.

39. Li H, Durbin R. 2009. Fast and accurate short read alignment with Burrows-Wheeler transform. Bioinformatics 25:1754–1760. DOI: 10.1093/bioinformatics/btp324.

40. Lubchenco J. 1995. The Role of Science in Formulating a Biodiversity Strategy. BioScience 45:S7–S9. DOI: 10.2307/1312437.

41. Luikart G, England PR, Tallmon D, Jordan S, Taberlet P. 2003. The power and promise of population genomics: from genotyping to genome typing. Nature Reviews Genetics 4:981–994. DOI: 10.1038/nrg1226.

42. Mardis ER. 2008. The impact of next-generation sequencing technology on genetics. Trends in Genetics 24:133–141. DOI: 10.1016/j.tig.2007.12.007.

43. McKenna A, Hanna M, Banks E, Sivachenko A, Cibulskis K, Kernytsky A, Garimella K, Altshuler D, Gabriel S, Daly M, DePristo MA. 2010. The genome analysis toolkit: A MapReduce framework for analyzing next-generation DNA sequencing data. Genome Research 20:1297–1303. DOI: 10.1101/gr.107524.110.

44. McKinney GJ, Larson WA, Seeb LW, Seeb JE. 2017. RADseq provides unprecedented insights into molecular ecology and evolutionary genetics: comment on Breaking RAD by Lowry et al . (2016). Molecular Ecology Resources 17:356-361. DOI: 10.1111/1755-0998.12649.

45. Meek MH, Larson WA. 2019. The future is now: Amplicon sequencing and sequence capture usher in the conservation genomics era. Molecular Ecology Resources 19:795–803. DOI: 10.1111/1755-0998.12998.

46. Meirmans PG. 2006. Using the AMOVA framework to estimate a standardized genetic differentiation measure. Evolution 60:2399–2402. DOI: 10.1111/j.0014-3820.2006.tb01874.x.

47. Milne I, Stephen G, Bayer M, Cock PJA, Pritchard L, Cardle L, Shaw PD, Marshall D. 2013. Using Tablet for visual exploration of second-generation sequencing data. Briefings in Bioinformatics 14:193–202. DOI: 10.1093/bib/bbs012.

48. Neethiraj R, Hornett EA, Hill JA, Wheat CW. 2017. Investigating the genomic basis of discrete phenotypes using a Pool0Seq0only approach: New insights into the genetics underlying colour variation in diverse taxa. Molecular Ecology 26:4990–5002. DOI: 10.1111/mec.14205.

49. O’Leary SJ, Puritz JB, Willis SC, Hollenbeck CM, Portnoy DS. 2018. These aren’t the loci you’e looking for: Principles of effective SNP filtering for molecular ecologists. Molecular Ecology 27:3193-3206. DOI: 10.1111/mec.14792.

50. Porras-Hurtado L, Ruiz Y, Santos C, Phillips C, Carracedo A, Lareu M V. 2013. An overview of STRUCTURE: Applications, parameter settings, and supporting software. Frontiers in Genetics 4. DOI: 10.3389/fgene.2013.00098.

51. Pritchard JK, Stephens M, Donnelly P. 2000. Inference of Population Structure Using Multilocus Genotype Data.

52. Rellstab C, Zoller S, Tedder A, Gugerli F, Fischer MC. 2013. Validation of SNP allele frequencies determined by pooled next-generation sequencing in natural populations of a non-model plant species. PLoS ONE 8. DOI: 10.1371/journal.pone.0080422.

53. Russello M, Amato G, DeSalle R, Knapp M. 2020. Conservation Genetics and Genomics. Genes 11:318. DOI: 10.3390/genes11030318.

54. Schlotterer C, Tobler R, Kofler R, Nolte V. 2014. Sequencing pools of individuals - mining genome-wide polymorphism data without big funding. Nature Reviews Genetics 15:749–763. DOI: 10.1038/nrg3803.

55. Schwarze K, Buchanan J, Taylor JC, Wordsworth S. 2018. Are whole-exome and whole-genome sequencing approaches cost-effective? A systematic review of the literature. Genetics in Medicine 20:1122–1130. DOI: 10.1038/gim.2017.247.

56. Shafer ABA, Peart CR, Tusso S, Maayan I, Brelsford A, Wheat CW, Wolf JBW. 2017. Bioinformatic processing of RAD0seq data dramatically impacts downstream population genetic inference. Methods in Ecology and Evolution 8:907–917. DOI: 10.1111/2041-210X.12700.

57. Sharma A, Lee JS, Dang CG, Sudrajad P, Kim HC, Yeon SH, Kang HS, Lee S-H. 2015. Stories and Challenges of Genome Wide Association Studies in Livestock - A Review. Asian-Australasian Journal of Animal Sciences 28:1371–1379. DOI: 10.5713/ajas.14.0715.

58. Sievert C. 2020. Interactive Web-Based Data Visualization with R, plotly, and shiny. Chapman and Hall/CRC.

59. Tange O. 2011. GNU Parallel: The Command-Line Power Tool. ;login: The USENIX Magazine 36:42–47. DOI: 10.5281/zenodo.16303.

60. Taus T, Futschik A, Schlotterer C. 2017. Quantifying Selection with Pool-Seq Time Series Data. Molecular Biology and Evolution 34:3023–3034. DOI: 10.1093/molbev/msx225.

61. Taylor HR, Dussex N, van Heezik Y. 2017. Bridging the conservation genetics gap by identifying barriers to implementation for conservation practitioners. Global Ecology and Conservation 10:231–242. DOI: 10.1016/j.gecco.2017.04.001.

62. Team Rs. 2020. RStudio: Integrated Development for R.

63. Teske PR, Golla TR, Sandoval-Castillo J, Emami-Khoyi A, van der Lingen CD, von der Heyden S, Chiazzari B, Jansen van Vuuren B, Beheregaray LB. 2018. Mitochondrial DNA is unsuitable to test for isolation by distance. Scientific Reports 8:8448. DOI: 10.1038/s41598-018-25138-9.

64. Toews DPL, Brelsford A. 2012. The biogeography of mitochondrial and nuclear discordance in animals. Molecular Ecology 21:3907–3930. DOI: 10.1111/j.1365-294X.2012.05664.x.

65. Toonen RJ, Puritz JB, Forsman ZH, Whitney JL, Fernandez-Silva I, Andrews KR, Bird CE. 2013. ezRAD: a simplified method for genomic genotyping in non-model organisms. PeerJ 1:e203. DOI: 10.7717/peerj.203.

66. Visscher PM, Wray NR, Zhang Q, Sklar P, McCarthy MI, Brown MA, Yang J. 2017. 10 Years of GWAS Discovery: Biology, Function, and Translation. The American Journal of Human Genetics 101:5–22. DOI: 10.1016/j.ajhg.2017.06.005.

67. Waldron A, Miller DC, Redding D, Mooers A, Kuhn TS, Nibbelink N, Roberts JT, Tobias JA, Gittleman JL. 2017. Reductions in global biodiversity loss predicted from conservation spending. Nature 551:364–367. DOI: 10.1038/nature24295.

68. Weir BS, Hill WG. 2002. Estimating F-statistics. Annu. Rev. Genet 36:721–50. DOI: 10.1146/annurev.genet.36.

69. Wepfer PH, Nakajima Y, Sutthacheep M, Radice VZ, Richards Z, Ang P, Terraneo T, Fujimura A, Toonen RJ, Mikheyev AS, Mitarai S, Economo EP. 2021. Inclusivity is key to progressing coral biodiversity research: Reply to comment by Bonito et al. 2021. Molecular Phylogenetics and Evolution:107135. DOI: 10.1016/j.ympev.2021.107135.

